# A transparent window into the rhizosphere: a simplified workflow for spatially resolved soil metabolomics

**DOI:** 10.64898/2026.02.13.705714

**Authors:** Harihar Jaishree Subrahmaniam, Paul Moritz, Phillip J. Becker, Maria Riedner, Simon Thomsen, Ina C. Meier, Kai Jensen

## Abstract

Root exudates play a central role in nutrient cycling, microbial recruitment, and plant-plant interactions, yet most experimental approaches for analyzing exudate chemistry rely on sterile hydroponic systems that poorly represent soil conditions. We present a low-cost, open-source, 3D-printed rhizobox platform and associated workflow that enable non-destructive root imaging and targeted rhizosphere soil sampling for LC-MS based metabolomics under realistic soil conditions. The design integrates a transparent removable window for repeated root observations, a defined soil volume to support spatially explicit sampling, and a blank-informed data-processing pipeline to distinguish plant-derived metabolites from soil and construction material background. We validated the system using the model plants *Arabidopsis thaliana* (Col-0) and *Phragmites australis*. We demonstrate reliable plant growth and consistent root development across the imaging window. We also show robust detection of species-specific rhizosphere metabolite profiles, with minimal variation in the vertical or temporal dimensions relative to the strong species effects. We further illustrate the application of the workflow in a factorial experiment manipulating social context (solo *vs*. conspecific pairs) and short-term heat stress in *A. thaliana*, showing that the approach is sensitive to treatment-associated changes in metabolite richness, diversity, and chemical composition in soil. The complete protocol, from rhizobox fabrication and assembly to soil extraction, LC-MS acquisition, and data curation can be implemented within 4-6 weeks using standard laboratory equipment and openly available design files. By combining ecological realism with analytical control, this workflow provides a broadly applicable method for quantifying rhizosphere metabolite dynamics across species, treatments, and spatial sampling zones, facilitating experimental studies of below-ground chemical processes in plant ecology.

## 1 Introduction

### 1.1 Root exudates as a methodological challenge in soil systems

Plants actively modify their below-ground environment through the release of a diverse array of low-molecular-weight compounds, including sugars, amino acids, organic acids, and secondary metabolites. These root-derived compounds contribute to nutrient mobilisation, microbial recruitment, and plant-plant interactions, linking plant physiology to soil ecological processes (Bais *et al*., 2006; Mommer *et al*., 2016; Canarini *et al*., 2019; Anten & Chen, 2021; Ma *et al*., 2022). Despite their recognised importance, experimental investigation of root exudate chemistry remains methodologically constrained, particularly under realistic soil conditions (Oburger & Jones, 2018; Casas & Matamoros, 2021). Root exudation occurs within an opaque, microbially active, and spatially heterogeneous matrix, where metabolites are rapidly transformed and diluted. Consequently, detection and interpretation of exudate-derived signals depend strongly on sampling position relative to roots and on the ability to distinguish plant-derived compounds from soil background (Ritter *et al*., 2025). Many existing approaches rely on destructive sampling that disrupts soil structure, obscures fine-scale spatial patterns, and ultimately limits reproducibility across replicates. Methods that enable consistent, spatially explicit rhizosphere sampling while preserving soil structure are therefore essential for linking chemical profiles to plant traits and experimental treatments over time.

### 1.2 Limitations of current systems for studying below-ground interactions

Experimental studies of plant-plant interactions increasingly recognize that below-ground chemical cues contribute to neighbor detection, kin recognition, and stress context-dependent responses (Dudley & File, 2007; Biedrzycki *et al*., 2010; Semchenko *et al*., 2014; Subrahmaniam *et al*., 2018). However, most empirical work in this area relies on hydroponic, agar-based, or split-root systems that offer analytical simplicity at the cost of ecological realism. Soil-based methods, including cuvette-based in situ exudate collection and whole-root dipping approaches, reduce this gap but still involve disturbance or short-term collection windows, constraining spatially explicit metabolomic analyses. While such systems are valuable for identifying candidate signalling compounds, they cannot capture the physical structure, diffusion constraints, and microbial activity characteristic of soil environments (Oburger & Jones, 2018; Ritter *et al*., 2025). Moreover, Petri plate and split-root designs constrain root growth to two-dimensional planes or artificially partition soil volumes, limiting their applicability to questions requiring spatially resolved sampling from rhizosphere zones occupied by specific roots or neighbors. As a result, it remains difficult to experimentally address where in the soil matrix chemical interactions occur and how metabolite profiles vary across neighboring root zones under controlled yet realistic conditions.

### 1.3 Abiotic stress and experimental complexity in soil metabolomics

Abiotic stressors such as heat and drought induce pronounced metabolic adjustments in plants, which often involve shifts toward osmoprotectants, antioxidants, and phenolic compounds (Bita & Gerats, 2013; Gargallo-Garriga *et al*., 2018). These changes can propagate into the rhizosphere, influencing microbial activity and chemical composition of soil adjacent to roots (Rolfe *et al*., 2019). Importantly, stress responses may depend on social context, with neighbor presence altering resource allocation or modulating chemical outputs through facilitative or competitive interactions (Vives-Peris *et al*., 2020; Pang *et al*., 2021; Sharma *et al*., 2023). Other below-ground interactions, such as those with mycorrhizal fungi or rhizoplane bacteria, can also influence root exudation patterns and rhizosphere chemistry. Despite growing interest in biotic × abiotic interactions below ground, very few experimental systems allow simultaneous manipulation of neighbour identity and environmental stress while remaining compatible with untargeted metabolomic analysis in soil (Ritter *et al*., 2025). This limitation reflects both practical constraints on spatial sampling and analytical challenges associated with background signal, material artefacts, and reproducibility in soil-based metabolomics.

### 1.4 Technical bottlenecks in soil-based rhizosphere metabolomics

Advances in metabolomic approaches have greatly expanded the scope of plant chemical profiling. Among these, liquid chromatography coupled to mass spectrometry (LC-MS) has become the most widely used technique for untargeted analysis of root-derived metabolites due to its high sensitivity and broad compound coverage. However, applying LC-MS based metabolomics to soil systems remains technically demanding. Hydroponic and agar-based approaches provide clean matrices and high analytical sensitivity, but they lack realistic diffusion gradients and microbial turnover (Oburger & Jones, 2018). In contrast, although soil-based experiments offer ecological relevance, they introduce a substantial background signal from soil organic matter, growth substrates, and experimental materials. A further challenge lies in spatial control. While many existing rhizobox designs prioritize root imaging, they were not developed for large-scale, replicated, and treatment-rich metabolomics experiments. They often lacking fixed sampling geometries (*i.e.,* predefined soil volumes and distances from roots), consistent sampling interfaces, and systematic blank controls, all of which are essential for scalable chemical analyses. Although recent hydroponic workflows have standardized exudate collection for metabolomics (Subrahmaniam *et al*., 2023; Döll *et al*., 2024), we are not aware of any widely adoptable, open-source systems that integrate root visibility, spatially defined soil sampling, and blank-informed metabolomic workflows.

### 1.5 A modular rhizobox approach for soil metabolomics

Rhizoboxes are growth containers with transparent windows for root observation that offer a promising compromise between analytical control and ecological realism. However, many available designs are costly, bulky, or suited to small number of species. To address these limitations, we developed a low-cost, open-source, 3D-printed rhizobox optimized for rhizosphere metabolomics in soil. The system incorporates a transparent, removable front window that enables repeated, non-destructive imaging of root systems and guides targeted rhizosphere soil sampling. Defined geometry and modular components support reproducible sampling positions across replicates and experimental treatments. The design is compatible with standard LC-MS extraction protocols and incorporates controls for both soil matrix and construction materials, enabling blank-informed filtering of background signals.

Hydroponic systems typically quantify metabolites released directly into the surrounding solution, providing a simplified representation of root exudation, but excluding soil structure, microbial turnover, and physicochemical retention processes. In contrast, soil-based sampling captures the pool of exudate-derived metabolites that persist, transform, or accumulate within the rhizosphere. These rhizosphere metabolites thus represent the chemically active environment experienced by neighboring roots and soil microbes and therefore provide a biologically meaningful proxy for below-ground chemical interactions under realistic conditions.

### 1.6 Study aims

In this study, we present and validate an experimental workflow for quantifying rhizosphere metabolite dynamics in soil using the 3D-printed rhizobox system. Specifically, our aims are:

1. Demonstrate that the rhizobox supports consistent plant establishment and root development under soil conditions, and that the workflow yields reproducible rhizosphere metabolite profiles that are clearly distinguishable from soil/material controls across biological replicates.
2. Assess cross-species applicability by comparing the rhizosphere metabolite profiles of the two model species *Arabidopsis thaliana* (Col-0) and *Phragmites australis*; and
3. Evaluate the sensitivity to biologically relevant treatments by conducting a factorial experiment that manipulates social context (solo *vs*. conspecific pairs) and short-term heat stress.

By detailing the design, sampling strategy, and analytical workflow, this study provides a first scalable and accessible framework for integrating metabolomics into experimental plant ecology. Our rhizobox setup enables mechanistic investigation of below-ground chemical processes under controlled yet realistic soil conditions.

## 2 Materials and Methods

### 2.1 Overview

We developed and validated a low-cost, open-source 3D-printed rhizobox workflow that integrates non-destructive root imaging with spatially explicit rhizosphere soil sampling and LC-MS based metabolomics under realistic soil conditions. Two experiments were conducted. Experiment 1 (April-July 2025) validated the performance of the rhizobox with the two model species *A. thaliana* (Col-0) and *P. australis*, which were cultivated under identical greenhouse conditions and sampled repeatedly over time and two vertical root zones. Experiment 2 (October-November 2025) applied the workflow to a factorial experiment, which manipulated the social context of the plants (solo *vs*. conspecific pairs) and air temperature (ambient *vs*. short term heat stress) in *A. thaliana* (Col-0). The latter included within-box, neighbor-resolved spatial sampling. The mass spectrometry data were processed using Mzmine3 (Schmid *et al*., 2023) and annotated with SIRIUS 5 (Dührkop *et al*., 2013). They were then analyzed statistically in*burg* (version 4.5.2) (R Core Team, 2025), using RStudio (version 2025.09.2+418) (Posit Team, 2025). Blank-informed filtering was used to separate the biological signal from the background noise by soil and materials.

### 2.2 Rhizobox design and assembly

Rhizoboxes were produced from a high-flow variant of polyethylene terephthalate glycol (PETG-HF) filament (Bambu Lab, Shenzhen, China), using a Bambu Lab X1E Fused Deposition Modeling (FDM) 3D printer (Bambu Lab, Shenzhen, China). All G-code was generated using Bambu Studio (v1.10.2.64, Bambu Lab). The rhizoboxes consisted of an opaque frame measuring 97 × 94 × 10 mm internally, with a perforated base to allow drainage. The transparent front window was made from a 2 mm-thick sheet of PLEXIGLAS® XT (extruded acrylic glass, PMMA, Röhm GmbH, Germany) and was attached using metal clamps. This enabled root imaging and access for exudate sampling. To exclude light, the window was fully covered with a removable 0.3 mm thick, opaque PVC sheet, with the edges sealed with matt black tape. Boxes were mounted vertically at 45° on a custom frame to direct root growth. All rhizobox design files, including printable STL files and a dimensioned schematic, are provided as Supplementary Data S1 and S2.

All components were rinsed with 96% ethanol and air-dried prior to use to remove surface residues; the system was not maintained sterile, allowing normal microbial recolonisation from soil. The inner surface of the Plexiglas front panel was coated with a thin layer of polytetrafluoroethylene (PTFE) spray to reduce soil adhesion and facilitate consistent rhizosphere sampling; PTFE is chemically inert and was confined to the window surface, minimising direct contact with roots or soil.

### 2.3 Soil preparation and greenhouse conditions

A standard peat-based soil (Einheitserde Type ED73; Einheitserde Werkverband, Germany) was sieved to remove coarse particles and homogenized. Rhizoboxes were loosely packed to a consistent fill height and moistened evenly with sterile distilled water prior to sealing the window. Throughout experiments, moisture was maintained by surface watering using a spray bottle to avoid waterlogging. No fertilizer was applied, as the aim was to maintain uniform, moderate nutrient conditions and avoid treatment-independent shifts in rhizosphere chemistry. The experiments were conducted in a climate-controlled greenhouse at the Universität Hamburg under the following conditions: air temperature 22 ± 1 °C, 16 h light / 8h dark photoperiod, and a photosynthetically active radiation (PAR) of 120–150 µmol m□² s□¹.

### 2.4 Plant cultivation

*A. thaliana* seeds were stratified for 72 h at 4 °C in darkness, germinated at 22 °C, and transplanted into rhizoboxes at the cotyledon stage using a moistened fine brush. *P. australis* was propagated in trays under greenhouse conditions and transferred into rhizoboxes at comparable early developmental stages to *A.thaliana*. All plants were grown in a greenhouse under a photoperiod of 16 hrs light and 8 hrs of darkness at an air temperature of 22 ± 1 °C.

### 2.5 Experimental designs and rhizosphere sampling

#### 2.5.1 Experiment 1: System validation for plant species, time intervals, and soil zones

To validate the rhizobox system for soil-based metabolomics, six rhizoboxes were established for each of two species (A. thaliana Col-0 and P. australis), together with three unplanted soil controls (15 rhizoboxes in total). Plants were grown for three weeks and sampled on three dates (13 June, 18 June, and 23 June 2025). At each sampling date, root systems were imaged by removing the transparent front panel. Immediately after imaging, rhizosphere soil was collected using sterile spatulas from within approximately 5 mm of visible roots to ensure close root proximity. Approximately 0.3 g of soil per sample was transferred to 2 mL tubes, kept on ice during processing, and stored at −80 °C until extraction. Sampling targeted two predefined vertical positions within each rhizobox: an upper soil zone (approximately 0–5 cm below the surface) and a lower soil zone (approximately 5–10 cm). To minimise disturbance artefacts from repeated sampling, soil was collected from different sides of the visible root system across time points.

#### 2.5.2 Experiment 2: Social context and heat treatment

To test the effects of social context and short-term heat stress on rhizosphere metabolomes, *A thaliana* (Col-0) was grown under three configurations: (i) solo (one plant per rhizobox), (ii) intraspecific pair (two plants per rhizobox, spaced approximately 2 cm apart), and (iii) unplanted soil controls. Seeds were sown in early October 2025 and transplanted into rhizoboxes at the cotyledon stage. Plants were grown under greenhouse conditions for approximately four weeks until the late vegetative rosette stage. Each configuration was exposed either to ambient conditions (22 °C) or to acute heat stress (45 °C for 2 h in a pre-heated chamber), followed by a 2 h recovery period at 22 °C. Five biological replicates were established per treatment × temperature combination (20 planted rhizoboxes total). All box positions were fully randomised within the greenhouse. Immediately after treatment, root systems were imaged and rhizosphere soil was collected using the same procedure as in Experiment 1. Samples were taken from within approximately 5 mm of visible roots, with approximately 0.3 g soil per sample transferred to 2 mL tubes, kept on ice, and stored at −80 °C until extraction. In pair treatments, sampling was spatially explicit: soil was collected from predefined neighbour-associated zones, including an inner interaction zone between roots and outer root-associated zones, allowing assessment of spatially resolved rhizosphere chemistry within individual rhizoboxes.

### 2.6 Phenotypic measurements and root imaging

Plants were photographed at each sampling date in Experiment 1 and once prior to treatment and sampling in Experiment 2. Aboveground traits (rosette diameter and leaf number) were measured using digital callipers. Roots were imaged by removing the transparent front panel and using a digital camera under greenhouse lighting. Root length and surface area were quantified in ImageJ/Fiji using the SmartRoot plugin (Lobet *et al*., 2011), with pixel-to-millimetre calibration based on a ruler included in each image.

### 2.7 Material and background controls

To account for non-biological background signals arising from soil matrix, extraction solvents, and rhizobox construction materials, multiple controls were included and processed in the same way as the planted samples. Because polymer surfaces and surface coatings can introduce chemical artefacts, material-specific controls were explicitly incorporated into both experiments. In Experiment 1, controls included unplanted soil to characterize soil-derived background, Plexiglas front panels with and without polytetrafluoroethylene (PTFE) coating as material controls, and solvent blanks to capture instrumental background. In Experiment 2, glass–soil controls were included as material controls to assess potential material- and heat-induced artefacts under elevated temperature conditions, alongside unplanted soil and solvent blanks. All controls were extracted, measured, and processed alongside the biological samples and were explicitly incorporated into the blank-informed filtering and limit-of-detection (LOD) definition during data processing (Section 2.11). These tests confirmed that neither the rhizobox material nor the PTFE coating showed detectable leaching under the conditions used.

### 2.9 Metabolite extraction

Metabolites were extracted from frozen soil following a modified protocol based on Pétriacq et al. (2017). Briefly, 0.2 g soil (*A. thaliana*) and 0.3 g soil (*P. australis*, Experiment 1), respectively, was mixed with 1 mL pre-cooled extraction solvent (methanol:water:formic acid, 80:19.9:0.1, v/v/v), vortexed for 30 s, and shaken horizontally for 45 min at 800 rpm and 4 °C. Samples were centrifuged at 14,000 × g for 15 min (4 °C), and supernatants were filtered through 0.22 µm PTFE filters into LC-MS vials and stored at −20 °C until analysis. All control samples were extracted in parallel using identical procedures. Because soil particles contain adsorption and ion-exchange sites, a fraction of root-derived metabolites may be retained or transformed within the soil matrix. The extraction therefore captures the pool of metabolites that are recoverable under the applied solvent conditions, representing the detectable rhizosphere metabolite fraction rather than the total amount of compounds released by roots.

### 2.10 LC-MS acquisition

LC-MS data were acquired on a Bruker maXis QTOF coupled to a Bruker Elute UHPLC. Chromatography was performed on a Waters BEH C18 column (2.1 × 100 mm, 1.8 µm) at 50 °C. Mobile phase A was 0.1% formic acid in water and mobile phase B was 0.1% formic acid in acetonitrile. Mobile phase B was held at 2 % for the first 3 min, afterwards, the gradient increased from 2% to 90% B over 10 min, was held for 1,5 min, and re-equilibrated at 2% B for 4 min (flow rate 0.5 mL min□¹). The injection volume was 5 µL per sample. The QTOF operated in positive Electrospray (ESI) mode (capillary voltage 4.5 kV; source temperature 200 °C; drying gas 9 L min□¹; scan range m/z 50–1500), with auto-MS/MS acquisition of the top five ions per cycle. Pooled QC samples were injected every ten runs to monitor retention-time and signal stability, and solvent blanks confirmed the absence of carry-over.

### 2.11 Data processing, blank-informed filtering, and annotation

#### 2.11.1 File conversion and feature detection

Raw vendor files (.d) were converted to the open mzML format using ProteoWizard (Chambers *et al*., 2012). Converted files were processed in MZmine 3 using identical parameter settings across all samples and experiments to ensure comparability. Processing steps included mass detection, chromatogram building, ADAP wavelet-based peak deconvolution, isotopic grouping, retention-time alignment, and gap filling (Supplementary Data S4). Feature tables and associated metadata were exported to R for downstream analysis. Signal intensities were treated as numeric values, with zero intensities retained to preserve true absence information. Data were log-transformed to stabilize variance and reduce the influence of extreme values, while maintaining sensitivity to low-abundance features. Where required for interpretation or summarization, linear intensities were reconstructed from transformed values.

#### 2.11.2 Blank-informed LOD filtering

Soil-based metabolomics is particularly susceptible to background signals originating from the soil matrix, extraction solvents, and experimental materials. To ensure that downstream analyses focused on biologically meaningful, plant-associated metabolites, we implemented a blank-informed feature filtering strategy that explicitly incorporated all relevant controls. Rather than applying a fixed global intensity cut-off, an adaptive, feature-specific limit-of-detection (LOD) approach was used to distinguish the biological signal from the background derived from soil, materials, and analytical? instruments. For each metabolite feature, the LOD was defined using only control samples as the median control intensity plus 2.5 times the median absolute deviation (MAD), providing a robust estimate of background variability. In Experiment 1, control samples included chemical blanks, soil controls, and material controls with and without polymer additives. Features were retained only if they showed no detectable signal in any control sample and exceeded their feature-specific LOD in at least three biological replicates of at least one species. This strict filtering ensured that retained features represented reproducible biological enrichment rather than background artefacts. In Experiment 2, LODs were calculated separately for ambient and heat-stress conditions to account for condition-dependent changes in background signal. Control samples matched to each condition were used where available. Features were required to exceed the condition-specific LOD in at least three biological replicates and to show a minimum three-fold enrichment relative to the maximum control signal. Features passing these criteria in at least one condition were retained for downstream analyses.

This adaptive filtering strategy balances sensitivity and specificity by minimizing false positives caused by soil or material artefacts while preserving low-abundance metabolites that reproducibly exceed background, resulting in a conservative, high-confidence feature set suitable for ecological interpretation.

#### 2.11.3 Compound annotation and pathway assignment

Putative compound annotation was performed using MS/MS-based analysis in SIRIUS 5. Molecular formulae were predicted from fragmentation trees, and candidate structures were inferred using CSI:FingerID. Chemical class assignments were obtained using CANOPUS / NPClassifier, enabling pathway-level interpretation even when exact structural identification was not possible. SIRIUS outputs were linked to MZmine features using explicit feature identifiers. In cases where multiple annotations mapped to a single feature, the annotation with the highest confidence and information content was retained. Curated annotations were used exclusively for class-level and pathway-based analyses.

### 2.12 Statistical analyses

All statistical analyses were conducted in R using established ecological and metabolomics workflows (Supplementary Data S3). For each sample, three complementary summary metrics were calculated: total signal intensity as a proxy for overall exudation, feature richness as a measure of chemical breadth, and Shannon diversity to capture both richness and evenness. For multivariate analyses, feature intensity matrices were log□-transformed prior to analysis. Multivariate patterns in metabolite composition were assessed using Bray-Curtis dissimilarities and principal component analysis (PCA), and visualized using metric multidimensional scaling (MDS). Permutational multivariate analysis of variance (PERMANOVA) was used to test the effects of species (Experiment 1) and social and thermal treatments (Experiment 2). Presence-absence based distance metrics were additionally used to confirm that observed patterns were not driven solely by differences in signal magnitude. Univariate responses were analyzed using linear models with false discovery rate (FDR) correction to account for multiple testing.

In Experiment 1, models tested species effects and included sampling date and vertical position. In Experiment 2, planned contrasts compared temperature effects within social treatments and social effects within temperature conditions. For pathway-level analyses, intensities were summed within chemical classes for each sample and converted to proportional class contributions. These class shares were analyzed using linear models appropriate to each experimental design, with *post hoc* contrasts performed using estimated marginal means and correction for multiple comparisons. Detailed parameter settings, annotation tables, and statistical outputs are provided in the Supplementary Information.

## 3 Results

### 3.1 System performance and LC-MS data quality across experiments

Across both experiments, plants established successfully in the 3D-printed rhizoboxes and developed root systems that consistently grew along the transparent front panel, enabling repeated non-destructive imaging and targeted rhizosphere soil sampling (Figure 1A, B). Neither experiment showed signs of waterlogging, severe wilting or systematic avoidance of the transparent surface, indicating that the rhizobox design provided stable growth conditions for both species and across treatments (Supplementary Table S1, S2). Soil moisture was monitored and maintained at similar levels across treatments, ensuring that the heat treatment represented primarily a temperature stress rather than a combined heat-and-drought treatment. Global ordinations of curated metabolite feature tables using MDS showed tight clustering of biological samples and pooled quality-control injections, with separation from soil, solvent, and material blanks in both experiments (Figure 1C, D). Importantly, different control types (soil-only, material, and solvent blanks) were found to cluster closely together and showed no systematic separation, indicating that the background signal composition was comparable across the controls. This separation indicates high analytical reproducibility and confirms that background signal derived from soil matrix, extraction solvents, and rhizobox materials can be effectively removed by the blank-informed preprocessing pipeline.

**Figure 1.**
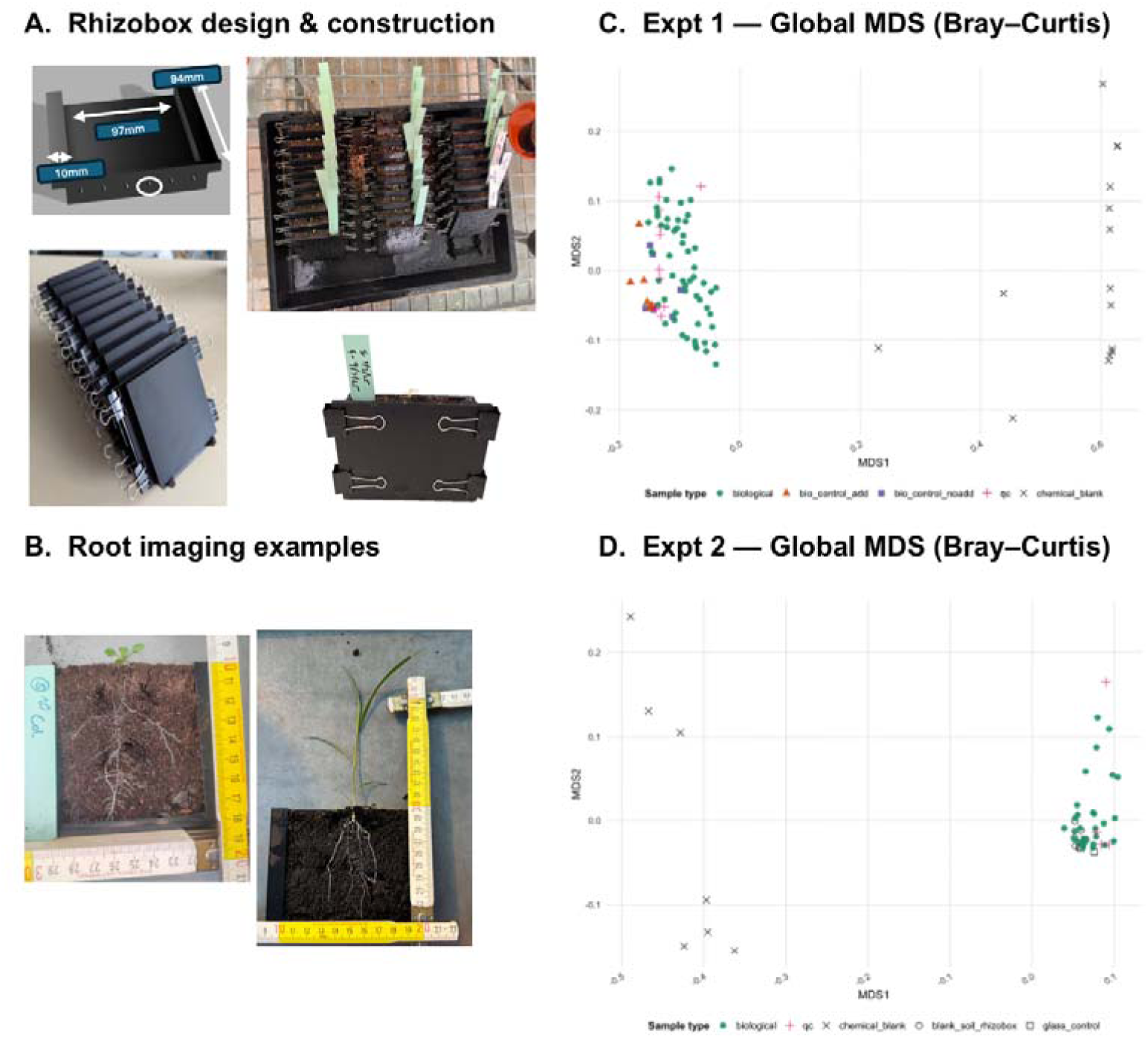
Rhizobox system and global structure of rhizosphere metabolomes. (A) Custom 3D-printed rhizobox design with removable plexiglass front for plant growth, root imaging, and rhizosphere sampling. (B) Representative *Arabidopsis thaliana* and *Phragmites australis* plants grown in rhizoboxes. (C) Global multidimensional scaling (MDS; Bray-Curtis dissimilarity) of rhizosphere metabolomes from Experiment 1, including biological samples, biological controls, quality control injections, and chemical blanks. (D) Global MDS (Bray-Curtis) of rhizosphere metabolomes from Experiment 2, showing samples from different social (solo *vs*. intra-specific) and thermal (ambient *vs*. heat) treatments alongside biological controls, quality control injections, and chemical blanks.

After the LOD sample-to-blank filtering, 24 features were retained from 1,358 detected features in Experiment 1, and 65 features were retained from 583 detected features in Experiment 2 (Supplementary Table S3, S4). In Experiment 2, a larger number of features exceeded condition-specific LOD thresholds under heat stress (52 features above LOD in heat *vs*. 27 in ambient, with 14 shared across temperature treatments), indicating increased detectability of rhizosphere metabolites under heat rather than systematic artefacts introduced by temperature or materials.

### 3.2 Species-specific rhizosphere metabolite profiles in Arabidopsis thaliana and Phragmites australis (Experiment 1)

Multivariate analyses revealed clear species-specific differences in rhizosphere metabolite composition. PCA of transformed feature intensities showed strong separation between A. thaliana and P. australis along the first principal component (Figure 2A). PERMANOVA on presence–absence profiles confirmed a significant effect of species identity (R² = 0.152, F = 7.87, p = 0.001), whereas sampling date (R² = 0.061, p = 0.111) and the species × date interaction were not significant (Supplementary Table S5). Univariate metrics supported these patterns. *A. thaliana* exhibited higher total rhizosphere signal (9492 ± 1338), feature richness (8.48 ± 0.80), and Shannon diversity (1.77 ± 0.10) than *P. australis* (2141 ± 739; 2.09 ± 0.38; 0.52 ± 0.11; Figure 2B–D, Supplementary Table S7). Feature-wise analyses further showed a strong main effect of species, with 16 of 24 curated metabolites differing significantly after FDR correction (Supplementary Table S9).

**Figure 2.**
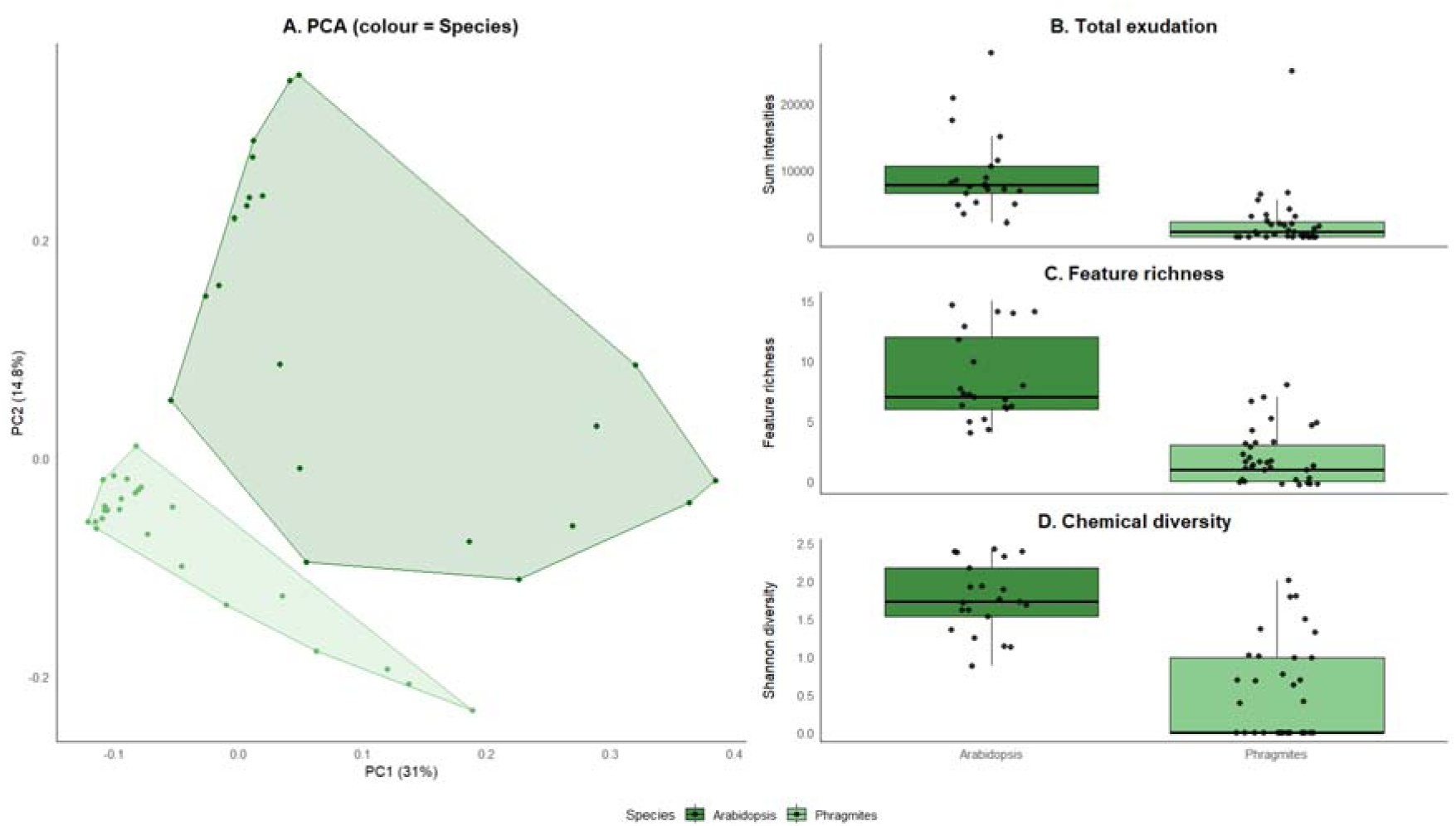
Multivariate structure and summary metrics for Experiment 1. (A) Principal component analysis (PCA) of log□-transformed rhizosphere metabolite feature intensities for *A. thaliana* and *P. australis*. (B) Total rhizosphere signal per sample, calculated as the sum of feature intensities. (C) Feature richness, defined as the number of detected metabolite features per sample. (D) Shannon diversity index calculated from relative feature intensities. Points represent individual samples.

Chemical class annotation indicated contributions from multiple rhizosphere-associated pathways, including fatty acids, amino-acid and peptide derivatives, alkaloids, polyketides, terpenoids, shikimate/phenylpropanoid derivatives, and unassigned compounds (Figure 3A). Class-level analyses revealed clear species contrasts, with *A. thaliana* enriched in amino-acid/peptide derivatives and fatty acids, and *P. australis* relatively enriched in secondary metabolite pathways. Temporal and vertical analyses showed little additional structuring. Class composition remained broadly stable across sampling dates (Figure 3C), and no features showed significant depth effects after FDR correction, indicating that species identity was the dominant driver of rhizosphere chemistry in this experiment.

**Figure 3.**
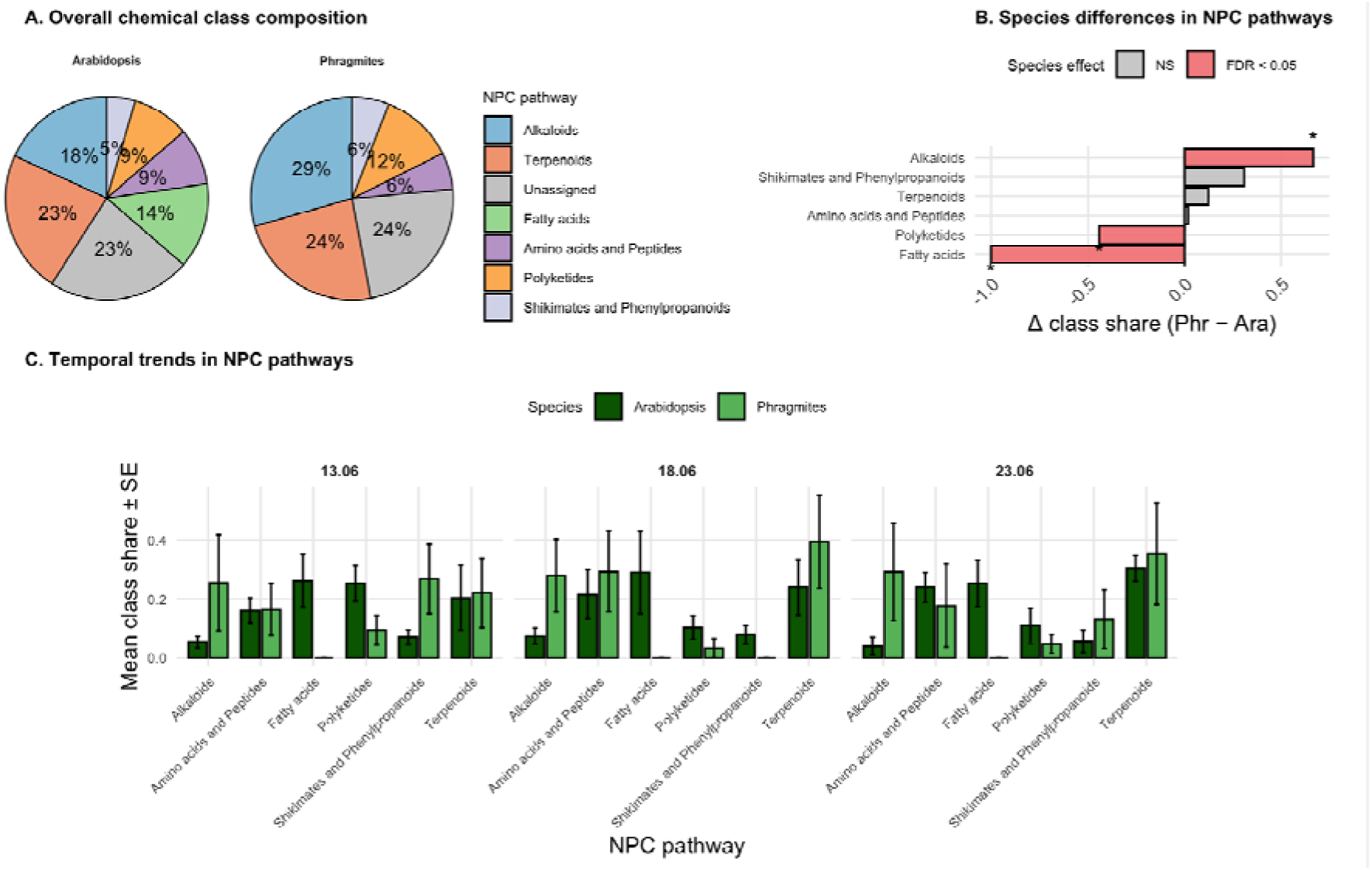
Chemical class composition and temporal dynamics of rhizosphere metabolomes in Experiment 1. (A) Overall chemical class composition of *A. thaliana* and *P. australis* rhizosphere metabolomes based on NPClassifier pathway annotations. (B) Differences in mean class share between species derived from two-way ANOVA results for NPC pathways. (C) Temporal variation in mean class share (± SE) across three sampling dates for major NPC chemical classes, shown separately for each species.

### 3.3 Effects of social context and heat stress on Arabidopsis rhizosphere metabolomes (Experiment 2)

Multivariate analyses of the curated dataset (65 features) revealed treatment-dependent structure in rhizosphere metabolite composition. A PERMANOVA on presence–absence profiles (Jaccard) including treatment, temperature, and their interaction explained 20.7% of total variation (R² = 0.207, F = 2.26, p = 0.008; Supplementary Table S6). PCA showed clear separation between solo and intra-specific pair treatments, whereas ambient and heat samples largely overlapped within each social context (Figure 4A). Term-level tests indicated no significant treatment × temperature interaction (R² = 0.009, F = 0.31, p = 0.937).

**Figure 4.**
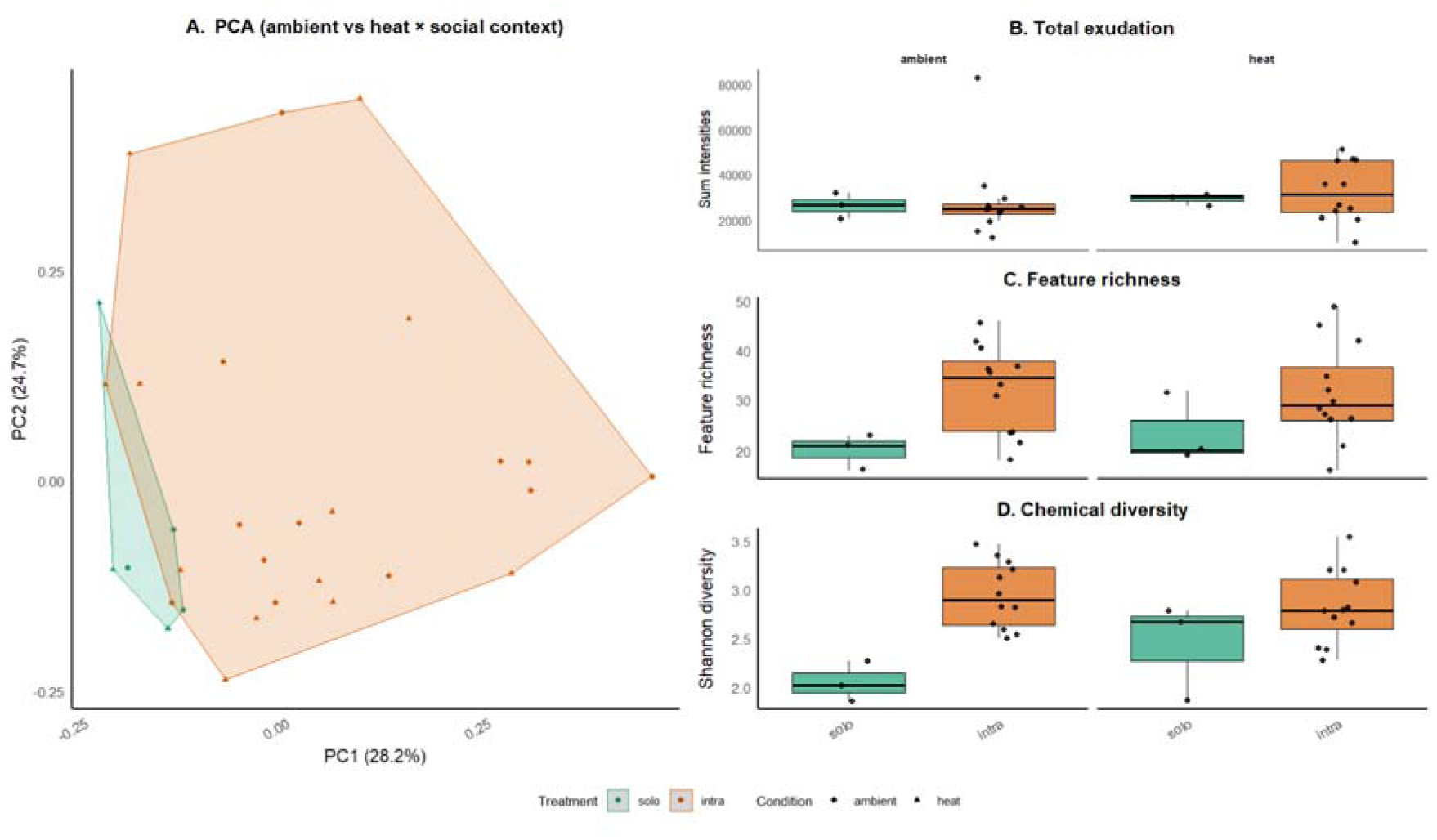
Effects of social context and temperature on *Arabidopsis thaliana* rhizosphere metabolomes (Experiment 2). (A) PCA of log□-transformed feature intensities across social (solo *vs.* intra-specific interaction) and thermal (ambient *vs*. heat) treatments (B) Total rhizosphere signal per sample across treatment combinations. (C) Feature richness across treatments. (D) Shannon diversity index across treatments.

Univariate metrics were consistent with these multivariate patterns (Figure 4B–D; Supplementary Table S8). Feature richness was higher in intra-specific pairs than in solo plants under both conditions (ambient: 32.5 ± 2.55 *vs.* 20.0 ± 2.08; heat: 31.4 ± 2.83 *vs.* 23.7 ± 4.18). Shannon diversity showed the same trend (ambient: 2.94 ± 0.10 *vs.* 2.05 ± 0.12; heat: 2.82 ± 0.11 *vs.* 2.44 ± 0.29 for intra *vs.* solo). In contrast, total signal varied only modestly across treatments, ranging from 26,330 ± 3,183 (solo ambient) to 32,569 ± 3,788 (intra heat) Supplementary Table S8). Feature-wise analyses detected few significant responses: across four planned contrasts (260 tests), only five comparisons remained significant after FDR correction (Supplementary Table S10). Chemical class annotation showed contributions from fatty acids, amino-acid and peptide derivatives, alkaloids, carbohydrates, polyketides, terpenoids, and an unassigned fraction (Figure 5A). Overall class composition differed only modestly between ambient and heat conditions, but temperature-associated shifts in class contributions varied between solo and intra-specific treatments (Figure 5B,C). Together, these results show that the rhizobox metabolomics workflow detects treatment-dependent structure in soil rhizosphere metabolomes, with social context exerting stronger effects than short-term heat stress.

**Figure 5.**
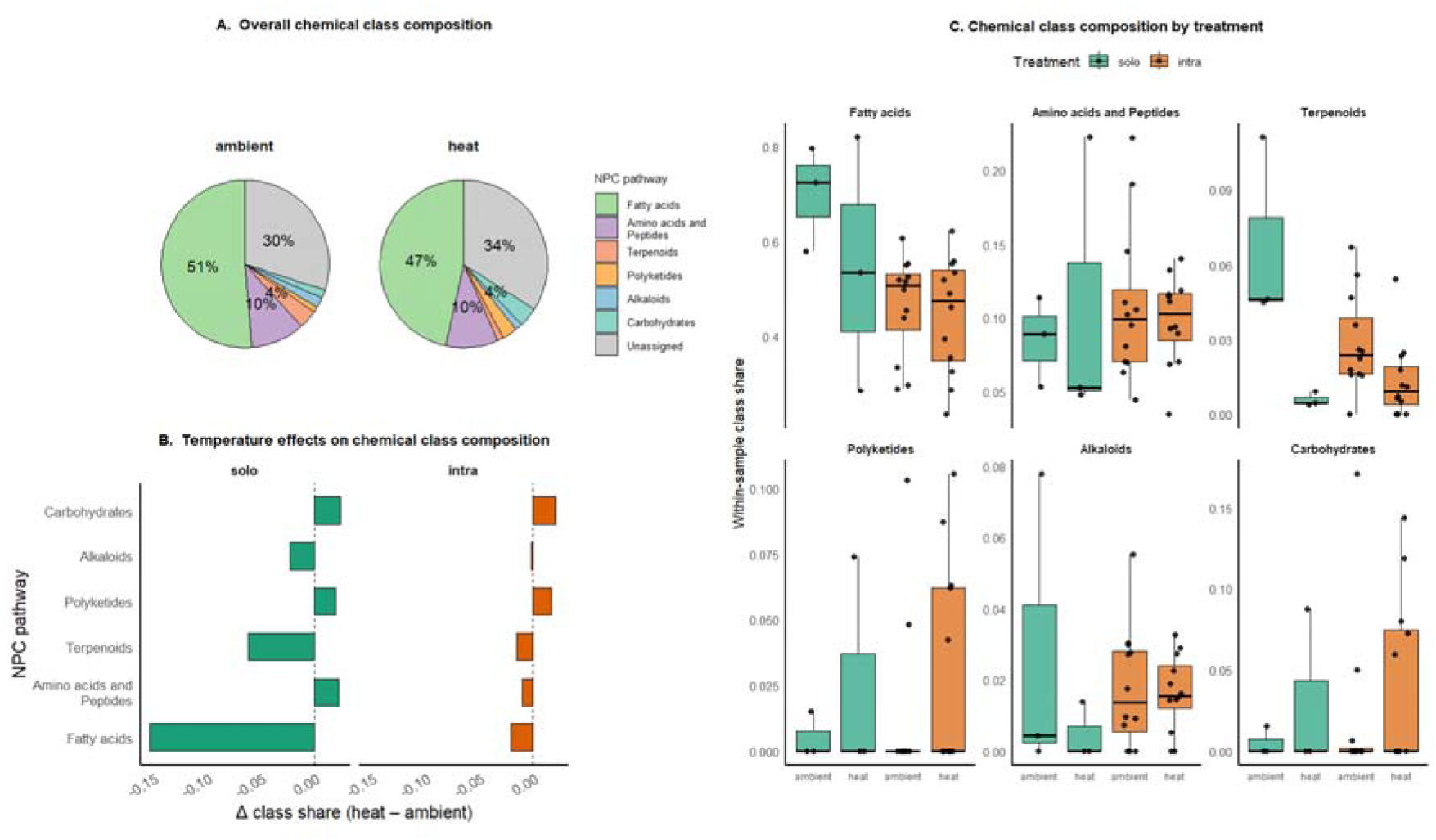
NPC chemical class structure of *A. thaliana* rhizosphere metabolomes under social and thermal treatments (Experiment 2). (A) Overall NPC chemical class composition under ambient and heat conditions. (B) Change in NPC class share between heat and ambient treatments, shown separately for social contexts. (C) Within-sample NPC class share across four combined combined social × thermal treatment groups.

## 4 Discussion

### 4.1 A modular platform for rhizosphere metabolomics in soil

This study presents a low-cost, 3D-printed rhizobox system for spatially resolved rhizosphere metabolomics under realistic soil conditions. Across two experiments, the system supported consistent plant growth, reproducible root development along the imaging surface, and reliable recovery of plant-associated metabolites from soil extracts. Ordination analyses clearly separated biological samples from blanks and material controls, indicating that plant-derived signals were detectable and could be distinguished from background profiles.

A key advantage of the system is the integration of non-destructive root imaging with targeted rhizosphere sampling. By visually confirming root presence before soil collection, the approach reduces spatial uncertainty compared to conventional pot experiments. At the same time, it retains soil structure and microbial activity, which are absent from hydroponic or agar-based systems. By combining soil realism with analytical control, the rhizobox provides an experimental bridge between molecular-scale metabolomics and genetic and ecological experimentation on root exudation, plant–plant interactions, and plant–microbe interactions. This framework enables metabolite-level measurements to be directly linked to plant genotypes, phenotypes, and interaction contexts under realistic soil conditions. Comparisons between soil-based rhizosphere metabolomes and hydroponic exudate profiles may further help disentangle compounds that are directly exuded from those shaped by soil-mediated transformation, offering a path toward integrating molecular, genetic, and ecological perspectives in future studies.

### 4.2 Performance across contrasting plant growth forms

Validation using *A. thaliana* and *P. australis* demonstrated that the rhizobox system accommodates species with markedly different root architectures and growth strategies. Despite differences in root morphology *i.e.,* fine and highly branched roots in *A. thaliana* versus coarser, clonal rhizomes in *P. australis*, both species established readily and produced consistent rhizosphere metabolite profiles across biological replicates. Accordingly, part of the higher total rhizosphere signal in *A. thaliana* likely reflects greater root development, in addition to species-specific differences in metabolite composition. This highlights the flexibility of the system and supports its applicability beyond small model plants.

Species-specific chemical signatures detected under identical soil and analytical conditions align with known physiological and ecological differences between these taxa. *A. thaliana* exhibited higher total signal, feature richness, and chemical diversity, with a greater proportional contribution of amino acids, peptides, and fatty acids, whereas *P. australis* showed relative enrichment of alkaloids, polyketides, and shikimate/phenylpropanoid-associated pathways. These patterns are consistent with previous reports linking *A. thaliana* exudation to low-molecular-weight, polar metabolites involved in nutrient acquisition, signalling, and microbial recruitment (Badri & Vivanco, 2009; Strehmel *et al*., 2014) and *P. australis* chemistry to phenolic compounds and secondary metabolites associated with allelopathy, oxidative rhizosphere modification, and clonal growth strategies (Armstrong *et al*., 1992; Bais *et al*., 2003; Rudrappa *et al*., 2007).

### 4.3 Social context as a dominant structuring factor of rhizosphere chemistry

Applying the rhizobox system to a factorial manipulation of social context and temperature revealed that conspecific neighbour presence was the primary driver of rhizosphere metabolome structure in *A. thaliana*. Multivariate analyses consistently separated solo and intra-specific pair treatments, whereas ambient and heat-stressed samples largely overlapped within each social context. Neighbour presence was also associated with higher feature richness and Shannon diversity, indicating that chemical heterogeneity in the rhizosphere increases when plants grow in close proximity to conspecifics.

Notably, total metabolite signal did not differ between solo and paired plants, suggesting that neighbour effects reflect redistribution of chemical investment across metabolites rather than increased overall release. At the level of individual features, relatively few metabolites responded strongly to social context after correction for multiple testing, consistent with the limited size of curated soil metabolite datasets. However, class-level analyses indicated that neighbour presence alters the proportional representation of several chemical pathways, supporting the view that social context influences the structure of the rhizosphere metabolome more strongly than any single compound. These findings align with a growing body of literature showing that plant-plant interactions influence root allocation, architecture, and physiology (Brooker *et al*., 2008; De Kroon *et al*., 2009; Novoplansky, 2009; Cahill *et al*., 2010; Subrahmaniam *et al*., 2018), but extend this work by demonstrating that social context leaves a detectable chemical signature in soil. Importantly, the rhizobox system enables such effects to be quantified while maintaining soil structure and microbial activity, which are essential for realistic chemical diffusion and transformation.

A methodological consideration in this experiment is that intra-specific pair treatments yielded a higher number of spatially resolved rhizosphere samples per rhizobox than solo plants, reflecting the neighbour-focused sampling design. While this increased spatial resolution strengthens inference about within-box chemical heterogeneity, overall biological replication was necessarily constrained by the labour-intensive nature of soil-based metabolomics. Importantly, this experiment was designed as a proof-of-concept to test whether social context leaves a detectable chemical signature in soil using a rhizobox-based metabolomics workflow.

### 4.4 Heat stress responses and their interaction with social context

Short-term heat stress induced detectable, but comparatively subtle, changes in rhizosphere chemistry relative to social context. Only a small number of individual metabolites responded significantly to heat after multiple-testing correction, including compounds annotated as lipid- and terpenoid-related metabolites with known roles in membrane stability and stress responses (Wahid *et al*., 2007; Bita & Gerats, 2013). At the chemical-class level, temperature effects manifested primarily as modest shifts in relative class contributions rather than wholesale reorganisation of the metabolome. Additionally, multivariate analyses did not support a strong interactive effect between heat stress and social context in this experiment. Instead, neighbour presence and temperature acted largely as additive or parallel influences, with social context consistently explaining more variation than temperature. This suggests that, under the stress regime applied here, heat alters specific components of the rhizosphere metabolite pool without overriding the broader structuring effect of neighbour presence.

These results caution against interpreting stress responses in isolation from biotic context, while also underscoring the importance of matching experimental claims to statistical support. The rhizobox system provides a framework for future studies that manipulate stress intensity, duration, and timing to test when and how abiotic stress reshapes chemically mediated plant interactions.

### 4.5 Rhizosphere metabolites as ecological intermediates

A key conceptual contribution of this work is the explicit focus on rhizosphere metabolites rather than root exudates sensu stricto. The compounds quantified here represent the fraction of plant-derived metabolites that persist, accumulate, or are transformed within the soil matrix, integrating plant release with microbial processing and physicochemical retention. As such, these metabolites constitute the chemical environment experienced by neighbouring roots and soil organisms, and may act as ecological intermediates, mediating interactions among co-occurring plants and between plants and microbes (Philippot *et al*., 2013; Jansson & Hofmockel, 2019). The species- and neighbour-specific patterns observed here suggest that plants structure their chemical surroundings in predictable ways, with potential consequences for subsequent root growth, microbial recruitment, or nutrient dynamics. Testing these mechanisms will require targeted experiments linking metabolite persistence to functional outcomes, but the rhizobox platform provides the spatial and temporal resolution needed to pursue such questions experimentally.

### 4.6 Methodological considerations and future directions

While the rhizobox system offers substantial advantages, several limitations should be acknowledged. The small soil volume constrains experiment duration and so far limits the amount of material available for parallel analyses, such as microbiome sequencing. Soil heterogeneity and microbial turnover may influence metabolite stability, and the acute heat-stress regime applied here captures short-term responses rather than longer-term acclimation. Future studies could extend this framework by incorporating time-series sampling, isotope labelling to trace metabolite origin and turnover, or combined metabolomic–microbiome approaches. The modular design also allows straightforward adaptation to multi-genotype combinations, sequential planting, or repeated stress exposures, enabling explicit tests of genotype-by-genotype and legacy effects mediated through soil chemistry.

### 4.7 Broader significance

By lowering the technical and financial barriers to soil-based metabolomics, the rhizobox protocol enables broader integration of chemical data into experimental plant ecology. It provides a practical means to test hypotheses concerning kin recognition, neighbour effects, and stress responses within an ecologically realistic framework. More broadly, the approach complements ongoing efforts to incorporate below-ground traits and processes into ecosystem and biodiversity models (Bardgett *et al*., 2014), where chemical interactions are increasingly recognised as key drivers of plant performance and community dynamics.

## 5 Conclusion

The 3D-printed rhizobox system enables reproducible cultivation, imaging, and spatially resolved rhizosphere sampling under realistic soil conditions. Validation with *Arabidopsis thaliana* and *Phragmites australis* demonstrated stable growth and robust recovery of species-specific rhizosphere metabolite profiles. Application to *A. thaliana* under contrasting social and thermal contexts showed that conspecific neighbour presence is a dominant driver of rhizosphere metabolome structure, while short-term heat stress induces more limited, pathway-specific effects. By making rhizosphere chemistry experimentally tractable without sacrificing soil realism, this protocol bridges molecular metabolomics and ecological experimentation. It provides a framework for mechanistic studies of how plants modify their chemical environment in response to neighbours and stress, advancing our understanding of below-ground interactions in plant communities.

## Acknowledgements

For both experiments, Col-0 seeds were obtained from the laboratory of Prof. Arp Schnittger (University of Hamburg), while *P. australis* plant material was kindly provided by Katrin Möller (University of Hamburg). We acknowledge the Technology Platform Mass Spectrometry of the University of Hamburg, University Medical Center Hamburg-Eppendorf and the Deutsche Forschungsgemeinschaft (DFG, German Research Foundation) for funding the mass spectrometer used in this work (project number 438807818). Ina C. Meier thanks the Deutsche Forschungsgemeinschaft (DFG, German Research Foundation) for financial support (Heisenberg program; grant no. ME 4156/5-1). We thank Birte Buske for her extensive support with chemical analyses and LC-MS measurements, as well as Annabelle Wang and Beatrice Money for their valuable assistance in conducting their bachelor’s thesis research within the framework of these experiments.

## Author contribution

H.J.S., I.C.M., and K.J. conceived and designed the study. S.T. designed and constructed the rhizobox system. P.M. and H.J.S. conducted the experiments. P.M., P.B., and H.J.S. performed chemical extractions. M.R. carried out LC-MS measurements. H.J.S. analyzed the data and wrote the first draft of the manuscript. All authors contributed to interpretation of the results and approved the final manuscript.

## Data Availability Statement

All data supporting the findings of this study are provided in the Supplementary Information and in public repositories. Supplementary Tables S1–S10 include sample metadata, curated metabolite feature tables, annotation results, multivariate statistics, and summary metabolome metrics for both experiments. Supplementary Data files include rhizobox design files, dimension schematics, analysis scripts, and detailed LC-MS data-processing parameters. All analysis scripts (R) and rhizobox design files (STL) are available via a public GitHub repository, with a permanent archival copy hosted on Zenodo. The Zenodo repository is available at: https://doi.org/10.5281/zenodo.18630194

## Supplementary Information

### Supplementary Tables

Table S1. Sample metadata for Experiment 1, including sampling date, species, zone, total root length (TRL) and total root surface area (RSA).

Table S2. Sample metadata for Experiment 2, including treatment, social context, temperature condition, TRL, and RSA.

Table S3. Curated metabolite feature table for Experiment 1 (n = 24 features after filtering).

Table S4. Curated metabolite feature table for Experiment 2 (n = 65 features after filtering).

Table S5. PERMANOVA and multivariate statistics summary for Experiment 1.

Table S6. PERMANOVA and multivariate statistics summary for Experiment 2.

Table S7. Summary metabolome metrics (total signal, feature richness, and Shannon diversity) for Experiment 1.

Table S8. Summary metabolome metrics (total signal, feature richness, and Shannon diversity) for Experiment 2.

Table S9. Feature-wise statistical tests for Experiment 1, including effect sizes and false discovery rate (FDR)-adjusted p-values.

Table S10. Feature-wise statistical tests for Experiment 2, including effect sizes and false discovery rate (FDR)-adjusted p-values.

### Supplementary Data

Data S1. Rhizobox 3D-print files (STL).

Data S2. Rhizobox dimension diagram (PDF).

Data S3. Analysis scripts (R) for Experiment 1 and Experiment 2.

Data S4. MZmine and SIRIUS processing parameters.

Data S5. Rhizosphere extraction protocol.

